# Phenotypic landscape of schizophrenia-associated genes defines candidates and their shared functions

**DOI:** 10.1101/360677

**Authors:** Summer B. Thyme, Lindsey M. Pieper, Eric H. Li, Shristi Pandey, Yiqun Wang, Nathan S. Morris, Carrie Sha, Joo Won Choi, Edward R. Soucy, Steve Zimmerman, Owen Randlett, Joel Greenwood, Steven A. McCarroll, Alexander F. Schier

**Author notes:** Correspondence may be addressed to Summer Thyme or Alexander F. Schier.

## Abstract

Genomic studies have identified hundreds of candidate genes near loci associated with risk for schizophrenia. To define candidates and their functions, we mutated zebrafish orthologues of 132 human schizophrenia-associated genes and created a phenotype atlas consisting of whole-brain activity maps, brain structural differences, and profiles of behavioral abnormalities. Phenotypes were diverse but specific, including altered forebrain development and decreased prepulse inhibition. Exploration of these datasets identified promising candidates in more than 10 gene-rich regions, including the magnesium transporter *cnnm2* and the translational repressor *gigyf2*, and revealed shared anatomical sites of activity differences, including the pallium, hypothalamus or tectum. Single-cell RNA sequencing uncovered an essential role for the understudied transcription factor *znf536* in the development of forebrain neurons implicated in social behavior and stress. This phenotypic landscape of schizophrenia-associated genes prioritizes more than 30 candidates for further study and provides hypotheses to bridge the divide between genetic association and biological mechanism.

## Introduction

Neuropsychiatric disorders are highly heritable. The understanding of their underlying genetic predispositions has recently accelerated, providing new opportunities to decipher disease mechanisms and discover potential drug targets (Pankevich et al., 2014). However, progress has been limited in part because the dozens of loci associated with these disorders contain hundreds of candidate genes (Grove et al., 2018; Schizophrenia Working Group of the Psychiatric Genomics, 2014; Wray et al., 2018). In particular, single nucleotide polymorphisms (SNPs) linked by genome-wide association study (GWAS) only define disease-associated genomic regions, which often span many genes (of which the culpable genes are unknown) or might include regulatory elements for distant genes (Won et al., 2016). Many of the same risk loci are also associated with multiple neuropsychiatric disorders (Cross-Disorder Group of the Psychiatric Genomics, 2013; Gandal et al., 2018; Grove et al., 2018; Wray et al., 2018) and even sleep disorders (Hammerschlag et al., 2017; Lane et al., 2017). This shared genetic predisposition indicates that some genes may contribute generally to brain health and development, while others may influence specific dimensions of disease manifestations (Cuthbert, 2014).

One of the richest of association datasets exists for schizophrenia (Schizophrenia Working Group of the Psychiatric Genomics, 2014). In 2014 the Schizophrenia Working Group of the Psychiatric Genomics Consortium identified 108 genomic loci at which common variants significantly associated with the disorder. Some identified genes align with ideas from previous research: for example, the implication of dopamine receptor D2 (*DRD2*, the target of many antipsychotic medications) may align with the dopamine hypothesis (Owen et al., 2016) or capture a pharmacogenetic effect on severity. The largest association signal, in the major histocompatibility complex (MHC) locus, appears to arise in part from the complement component 4 genes, which might contribute to excessive synaptic pruning (Feinberg, 1982; Glantz and Lewis, 2000; Sekar et al., 2016). However, many associated genes have never been studied, particularly in the context of nervous system development and activity.

To ascertain the mechanisms underlying polygenic illnesses such as schizophrenia and other neuropsychiatric disease, it would be useful to identify the *in vivo* functions of many associated genes; such analysis could prioritize candidate genes for further analyses. We reasoned that zebrafish, a small and low-cost vertebrate (Hoffman et al., 2016; Rennekamp and Peterson, 2015; Rihel et al., 2010), could be used to analyze orthologues of associated genes and that comprehensive mutant phenotyping could identify relevant candidates and suggest their impacts on brain function (Figure 1A). Previous studies have shown that zebrafish are a powerful model system to uncover developmental and behavioral functions of human genes (Escamilla, 2017; Haesemeyer and Schier, 2015; Hoffman et al., 2016). The two organisms share the majority of genes and tissues, including homologous brain structures (Wilson et al., 2002), and mutant screens have defined the functions of hundreds of conserved genes (Driever et al., 1996; Haffter et al., 1996; Kettleborough et al., 2013). Although human neuropsychiatric illness cannot be phenocopied in zebrafish, and homozygous knockouts differ from the functionally subtle polymorphisms that commonly segregate in human populations, neuroanatomical and behavioral phenotypes relevant to human biology can be scored in zebrafish (Burgess and Granato, 2007b; Meincke et al., 2004; Owen et al., 2016; Rihel et al., 2010; Wilson et al., 2002). Moreover, mutant analyses in zebrafish have yielded important insights into pathways underlying human neuropsychiatric illness (Escamilla, 2017; Hoffman et al., 2016).

**Figure 1.**
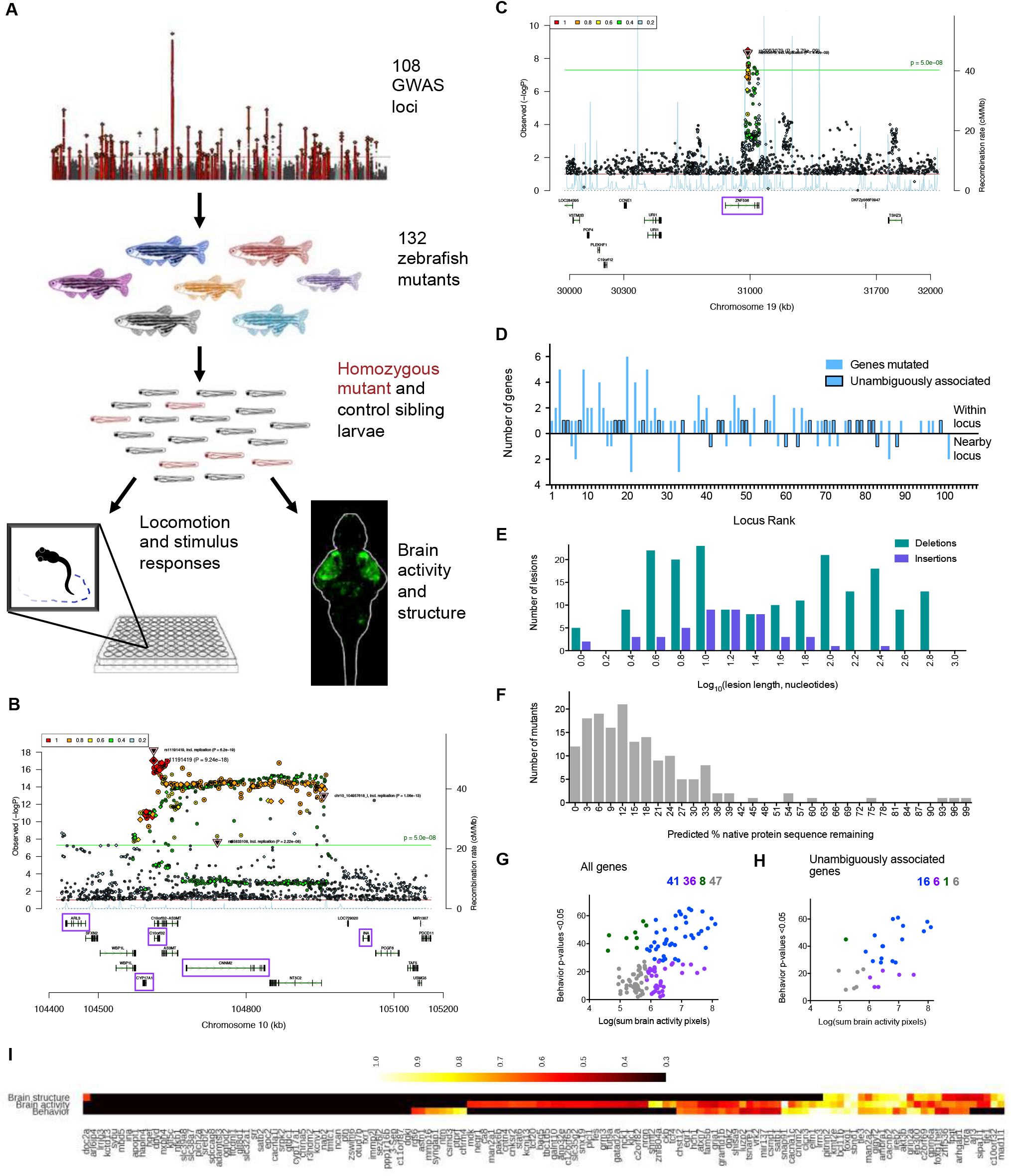
Generation and analysis of 132 mutants for schizophrenia-associated genes. (**A**) Mutants were generated via Cas9 mutagenesis for genes found within and neighboring genomic loci linked to schizophrenia through genome-wide association (Schizophrenia Working Group of the Psychiatric Genomics, 2014). Manhattan plot image was adapted from (Schizophrenia Working Group of the Psychiatric Genomics, 2014). The resulting mutants were assessed for changes to brain morphology, brain activity, and behavior. Multiple zebrafish orthologues existed for 30 of the 132 genes (162 individual zebrafish genes), and both copies were mutated and assessed together as one gene. (**B**) Ricopili plot (https://data.broadinstitute.org/mpg/ricopili/) for multi-gene locus #3 of 108 from which we selected five candidates (purple boxes) to mutate. (**C**) Ricopili plot for a gene (*znf536*) considered unambiguous because there are no other genes within 0.5 megabase on each side of the associated region, and the gene within the association is brain-expressed. (**D**) Mutants made from 79 of 108 associated genomic loci. The locus rank reflects the statistical strength of the genetic association (Schizophrenia Working Group of the Psychiatric Genomics, 2014), with 1 being the most significantly associated. A region of 0.5-2 MB around each locus was analyzed, and genes outside of the region of linkage disequilibrium were selected for 19 of the 79 loci. Unambiguously associated genes are implicated strongly by previous literature, such as genes involved in glutamatergic neurotransmission, or are the only genes within or neighboring their locus (Table S1). (**E**) Mutations generated from Cas9 cleavage. A range of mutations was recovered, tending to be either small (<15 bases) when a single gRNA cleaved or large (>100 bases) when a deletion spanned target sites of multiple gRNAs. Some mutants contained several lesions if multiple gRNAs cleaved the genome independently, and all are included here. (**F**) Amount of the protein predicted to remain in mutants, based on sequence alignment identity. This analysis included both orthologues if the gene was duplicated, for a total of 162 individual genes. The four mutants with >75% of the protein remaining did not have frameshifting mutations but did have phenotypes (Table S2), indicating that the protein function was disrupted. (**G**) Phenotypes in all 132 mutants based on analysis of brain activity signals and 71 behavioral assessments. See also Figure S1 for the cutoffs for classifying which mutants have phenotypes. (**H**) Phenotypes in mutants for the 29 unambiguously associated genes (Table S1). (**I**) Phenotype dimensions affected in mutants for 132 genes from 79 schizophrenia-associated loci. Quantification of brain activity, brain structure, and behavioral differences for mutants designated as having a phenotype (Figure S1, Figure 1G, Figure 3B, Table S2) was scaled for comparison between the three measures, with the weakest phenotype designated as 0.5 and strongest as 1.

To identify neurobiological functions and define candidate genes, we created mutants for 132 schizophrenia-associated genes and performed large-scale phenotypic analyses of brain activity, morphology and behavior. We established phenotypic databases, websites to share these datasets (genepile.com/scz_gwas108 and stackjoint.com/zbrain) (Supplementary Video 1), and robust analysis tools to facilitate large-scale studies. Guided by this dataset, we identified promising candidates in multi-gene loci and discovered neurobiological roles for more than 30 genes, including several that were largely understudied.

## Results

### Generation of 132 mutants for schizophrenia-associated genes

We focused on the 108 genomic loci at which common variants exhibited genome-wide significant associations in the largest schizophrenia case/control analysis (Schizophrenia Working Group of the Psychiatric Genomics, 2014), choosing 132 genes to mutate and phenotype in zebrafish (Figure 1A, Table S1, Table S2). For gene-rich loci, numerous candidate genes were individually mutated (Figure 1B). Criteria for choosing genes to mutate included nervous system expression (Lein et al., 2007) and human data that further implicated them in neuropsychiatric illness, such as patient exome sequencing or postmortem brain analyses of gene expression or chromatin conformation (Fromer et al., 2016; Won et al., 2016). While some candidates had known neurobiological functions, we mutated many minimally studied genes based only on nervous system expression (Lein et al., 2007). Twenty-nine of the 132 genes were strongly implicated by multiple lines of previous research or were the only candidate within and neighboring their respective locus (Figure 1C), and therefore were considered “unambiguously associated” with schizophrenia (Figure 1D, Table S1). These included well-known genes such as calcium channel subunits and the NMDA receptor subunit *grin2a*, minimally characterized brain-expressed candidates from single-gene loci such as *znf536* (Figure 1C) and *znf804a*, and genes implicated by CommonMind postmortem patient brain expression data (Fromer et al., 2016) such as *clcn3* and *snap91*. Selected genes were not always closest to the association signal. For example, at locus 83 we mutated *foxg1*, although it is almost 1 megabase from the associated region, instead of the closer *prkd1* because 1) the region overlaps with a known *FOXG1* enhancer (Allou et al., 2012; Ellaway et al., 2013) active in human neural progenitors (Won et al., 2016), 2) *foxg1* regulates forebrain development (Eagleson et al., 2007; Roth et al., 2010), and 3) this factor is involved in multiple neurodevelopmental disorders (Ariani et al., 2008; Mariani et al., 2015).

We used Cas9 and multiple guide RNAs (gRNAs) to mutagenize a conserved part of each protein near the N-terminus, predicted to result in substantial truncations (Figure 1E, Figure 1F, genepile.com/scz_gwas108). Only two homozygous mutants, *cacna1c* and *slc32a1* (Seisenberger et al., 2000; Stainier et al., 1996; Wojcik et al., 2006), were lethal prior to 6 days post-fertilization (dpf). An additional twenty-four zebrafish mutants (18%) appeared healthy at 6 dpf, but were lethal or underdeveloped by adulthood (Table S2). More than half of all mutants had phenotypes in brain activity or behavior (Figure 1G, Table S2, Figure S1), including more than 75% of those for unambiguously associated genes (Figure 1H). The presence and severity of brain activity and behavioral phenotypes were often but not always correlated (Figure 1G, Figure 1I).

### Baseline behavior and sensory stimulation

To uncover both baseline and stimulus-driven behavioral abnormalities, we subjected over 15,000 larvae (Table S2) to a range of behavioral paradigms from 4-6 dpf. We assessed baseline motion over multiple day-night cycles and responses to sensory stimulation (Burgess and Granato, 2007b; Kokel et al., 2010; Rihel et al., 2010). Phenotypic assays compared homozygous mutants to heterozygous and/or wild-type siblings and all larvae were genotyped. As it was unknown how perturbing these genes would manifest in larval zebrafish behavior, we developed software to measure a comprehensive range of parameters for both baseline movement and stimulus responses. We determined the frequency of movement (Figure 2A), features of movement (e.g. velocity, distance traveled) (Figure 2B), and location preference within the well of the multi-well plate (Figure 2C). These baseline parameters were calculated for 14 different time windows over the two-day experiments (Figure 2A, Figure 2E) to distinguish phenotypes that change over time such as sleep behavior. Sensory stimulation included response to light flash (Burgess and Granato, 2007a), dark flash (Wolman et al., 2011), and acoustic tap, as well as habituation to taps (Wolman et al., 2011), prepulse inhibition (Burgess and Granato, 2007b), and movement during a period of stressful heat. For startle responses to a stimulation event, such as a dark flash, multiple parameters of the response motion (e.g. speed, displacement) were calculated in addition to frequency of the response (Figure 2D).

**Figure 2.**
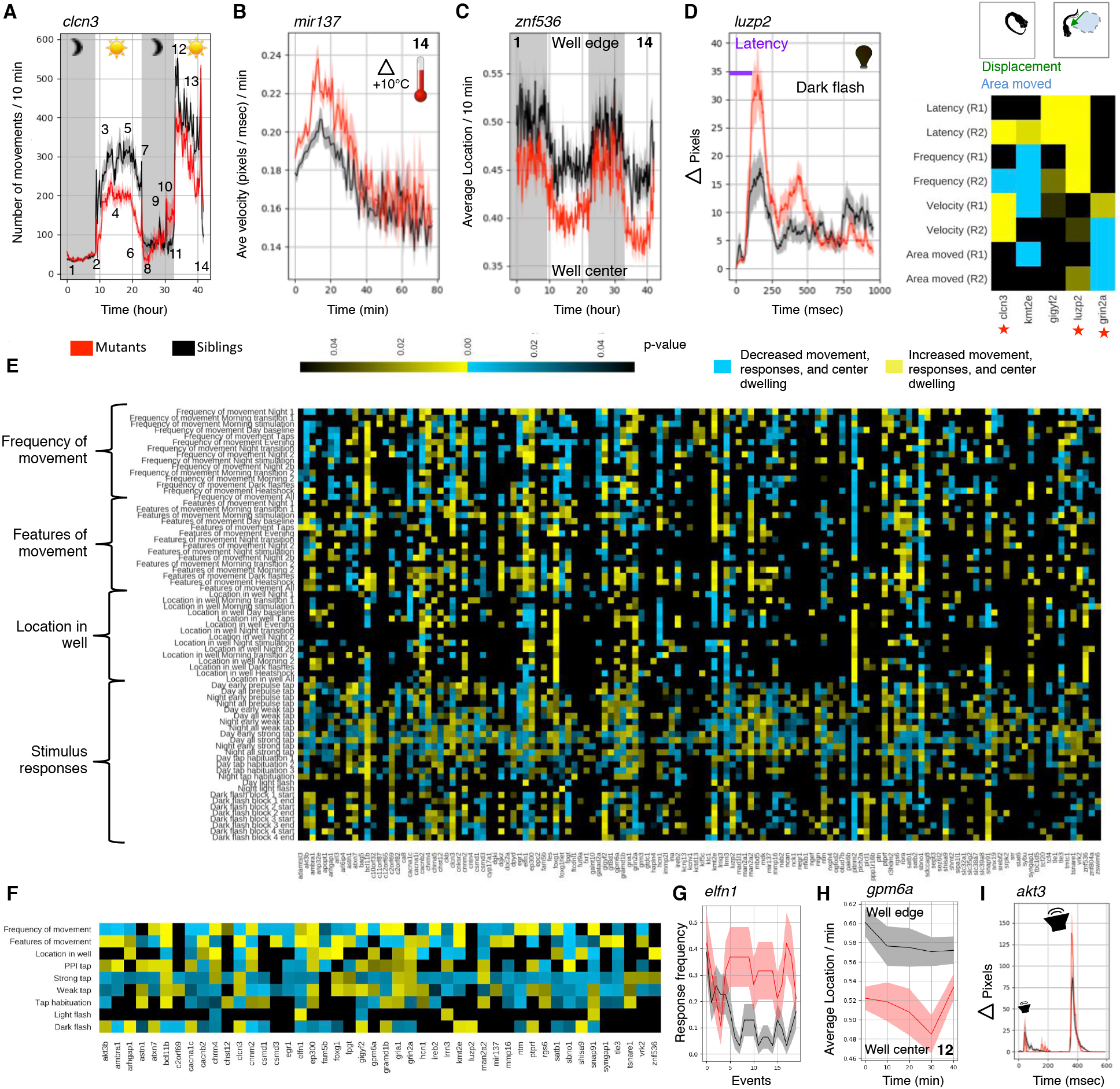
Behavioral phenotypes in zebrafish mutants. (**A**) Behavior protocol, displayed with the *clcn3* −/− mutants compared to +/− sibling controls and a frequency of movement metric. Frequency of movement, features of movement, and location preferences were calculated for each of 14 windows of the data: (1) Night 1, 9 hours, (2) Morning transition 1, 10 minutes, (3) Morning light flash, 125 minutes, (4) Day baseline, 165 minutes, (5) Day taps, including both tap habituation, prepulse tests, and tap response threshold tests, 5 hours and 22 minutes, (6) Evening, 217 minutes, (7) Night transition, 10 minutes, (8) Night 2, 110 minutes, (9) Night stimulation, including tap habituation, prepulse tests, tap response threshold tests, and light flash stimulation, 6 hours, (10) Night 2b, second section, 119 minutes, (11) Morning transition 2, 10 minutes, (12) Morning 2, 50 minutes, (13) Dark flashes, 7 hours, and (14) Heat stressor, >70 minutes. Behavioral tests were repeated for 73 of the mutants, 46 of those designated as having a phenotype. (**B**) Example of altered features of movement. (**C**) Example of altered location preference. (**D**) Example of a stimulus-driven high-speed response to a one second long dark flash, and quantification of dark flash phenotypes across various modalities of the response. The response graph is an average of larvae in the mutant and control groups for events where a response was observed. Dark flash responses were analyzed for four hour-long blocks of dark flashes by assessing the first ten and last ten flashes each block, totaling eight separate statistical analyses. These eight p-values were combined using fisher’s method to generate the heatmap of the five mutant phenotypes. (**E**) Summary of all 71 behavior assays for all tested mutants. If a mutant was repeated, the lowest combined p-value is shown here. If multiple comparisons were made for a given gene, such as in the case of duplicated genes, the lowest p-value comparison is shown. (**F**) The 46 mutants shown here had a total behavioral phenotype score greater than the cutoff (Figure S1) and were tested more than once. If the merged p-value (Figure S2) for the assay was significant (p < 0.05) between two independent repeats, the signal was counted and the lowest p-value of the two is displayed. Otherwise neither p-value was included. The significant signal had to be in the same time window or in the same stimulus assay to be counted (i.e., a significant phenotype observed on night 1 in one repeat and night 2 in another was not counted). See also Figure S2. (**G**) The *elfn1* mutants displayed increased responses to light stimulation. (**H**) The *gpm6a* mutants displayed a preference for the well center. (**I**) The *akt3* mutants displayed reduced prepulse inhibition. The mutant response profile demonstrates the prepulse behavior paradigm and response, where a weak tap (prepulse) is followed 300 milliseconds later by a strong tap. Plots of mutant compared to control groups in all panels in this figure represent mean ± s.e.m.. See also Figure S3.

We discovered mutants with phenotypes in each behavioral paradigm (Figure 2E). Many mutants were affected in multiple assays (Figure 2E, Figure 2F), although there were exceptions such as the highly specific increased dark flash response of mutants for the uncharacterized and unambiguously associated leucine zipper protein *luzp2* (Figure 2D, Figure 2F). Our fine-grained analysis method identified mutants with abnormalities in specific modalities of stimulus response (Figure 2D). Similarly, some mutants had subtle differences in their baseline motion, such as the increased velocity of movements in *mir137* mutants that was accentuated by heat stress (Figure 2B, Figure 2E). Other distinct phenotypes included increased light responsiveness in *elfn1* mutants (Figure 2G), decreased prepulse inhibition in *atxn7* mutants (Figure 2F), and a strong preference for the well center in *znf536* mutants (Figure 2C). Behavioral differences in zebrafish mutants also mirrored mammalian behavioral phenotypes. For example, the *gpm6a* mutant zebrafish preferred the well center just as *Gpm6a* mutant mice avoid closed spaces (El-Kordi et al., 2013) (Figure 2H), and mutants for *akt3* displayed decreased prepulse inhibition in zebrafish (Figure 2I) and mouse models (Bergeron et al., 2017). These *in vivo* datasets reveal the immense diversity of behavioral changes caused by mutations in schizophrenia-associated genes and provide entry points to study the underlying neural circuits.

### Whole-brain activity and morphology

We imaged more than 10,000 6 dpf larvae brains for altered brain activity and morphology in the 132 mutants (Table S2). Phenotypic assays compared homozygous mutants to heterozygous and/or wild-type siblings, and all larvae were genotyped. Whole-brain activity of unstimulated freely swimming larvae was determined by measuring the levels and distribution of phospho-Erk (Randlett et al., 2015), a downstream reporter of calcium signaling. Imaged brains were registered to the Z-Brain atlas and significant changes in activity between mutant and control groups were calculated (Randlett et al., 2015). Changes in brain volume in zebrafish mutants were calculated using deformation-based morphometry (Rohlfing and Maurer, 2003). All brain activity and structure maps can be explored in our website resource stackjoint.com/zbrain, where users can also upload and share their own imaging data (Supplementary Video 1).

Only sixteen (12%) of the mutants had substantial brain morphological differences (Figure 3A, Figure 3B, Table S2), compared to over half with brain activity differences (Figure 1G). Mutants for the transcription factors *foxg1* and *bcl11b* (Ctip2) had smaller forebrains (Figure 3C), consistent with their known forebrain development roles in mouse (Chen et al., 2008; Eagleson et al., 2007). Similarly, *rora* transcription factor mutants had underdeveloped cerebella (Dussault et al., 1998; Sidman et al., 1962), and *akt3* mutants had overall smaller brains (Easton et al., 2005). Thus, mutant brain anatomy differences agreed with previously described mammalian models. New neurodevelopmental roles were uncovered for lesser-studied genes such as the translational repressor *gigyf2* (Guella et al., 2011; Kryszke et al., 2016; Morita et al., 2012) and transcription factor *znf536*, which both had decreased volume in specific brain subregions including the forebrain (Figure 3C), and histone methyltransferase *kmt2e* (Ali et al., 2013) (Table S2, stackjoint.com/zbrain), which had increased brain volume (Figure 3C).

**Figure 3.**
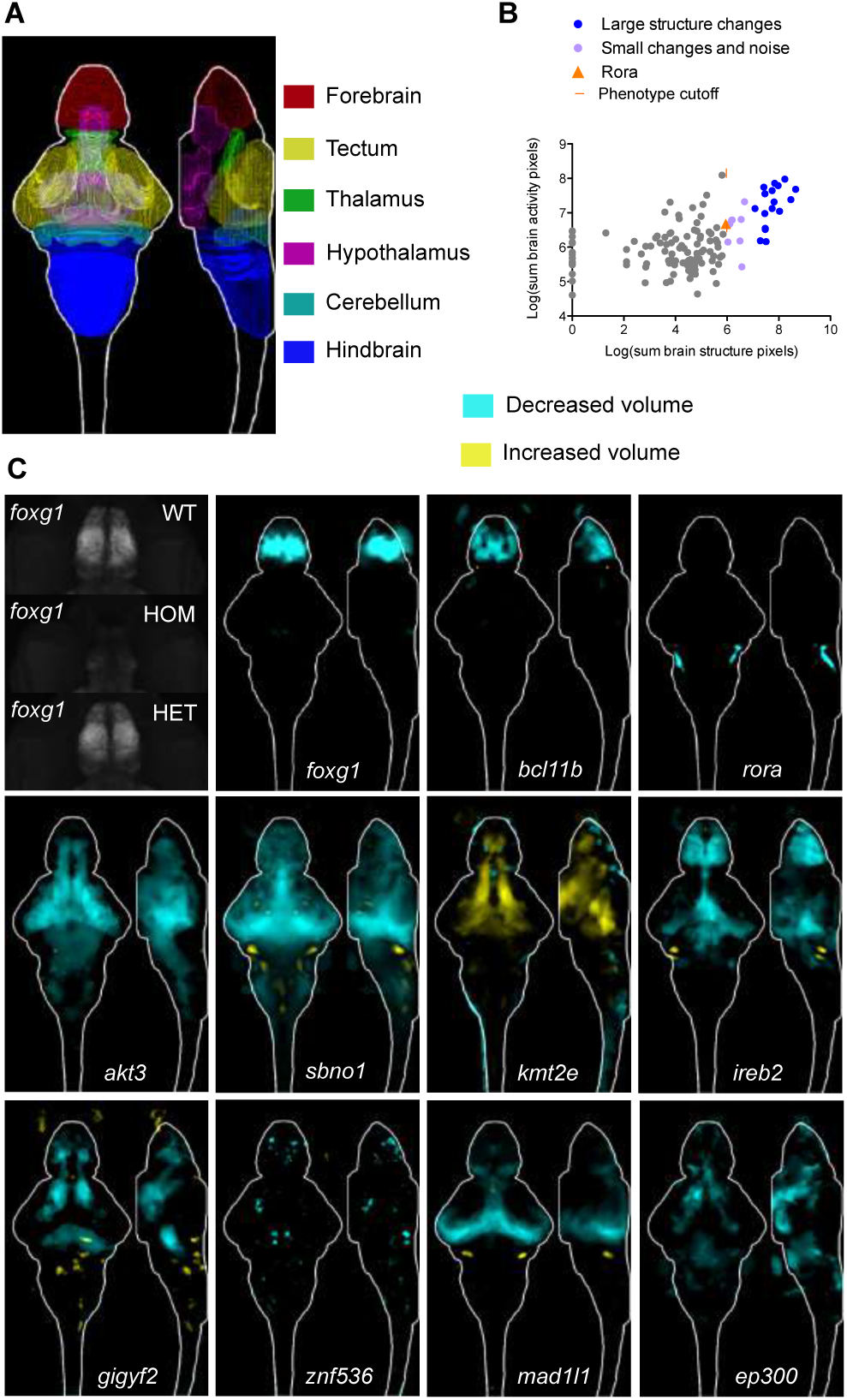
Whole-brain morphology phenotypes in zebrafish mutants. (**A**) Location of major regions in the zebrafish brain. (**B**) Comparison between brain activity and morphology data for all mutants. The *rora* mutant (orange triangle) represents the smallest (5.95) structural change designated as a phenotype, shown panel C. (**C**) Examples of structural differences in mutants calculated using deformation-based morphometry and displayed as sum-of-slices projections. Raw imaging data examples are shown for the *foxg1* mutants, demonstrating that the forebrain in homozygous (HOM) mutants is severely underdeveloped. A reduction in forebrain size of heterozygous (HET) *foxg1* mutants when compared to wild type (WT) siblings can be quantified (sum-of-slices projection), although it is not readily apparent in the raw data.

More than half of the mutants had brain activity phenotypes (Figure 1G, Figure 4), including 75% of genes unambiguously associated with schizophrenia (Figure 1H, Table S1). Decreased and increased activity differences emerged in both broad and localized patterns (Figure 4A, Figure 4B, Table S2, stackjoint.com/zbrain). For example, the unambiguously associated complement pathway regulator *csmd1* (Escudero-Esparza et al., 2013) exhibited broad activity increases. Elevated complement component 4 levels are observed in schizophrenia patients and may contribute to aberrant synaptic pruning (Sekar et al., 2016). In contrast, mutants for two unambiguously associated ion channel genes, *cacna1c* (Stainier et al., 1996) and *clcn3*, had localized forebrain activity decreases. Mutant brain activity phenotypes did not always spatially correlate with gene expression patterns (Figure S4, stackjoint.com/basic), as exemplified by the widespread expression of *clcn3* and its localized brain activity phenotype. Although the molecular pathways underlying schizophrenia are unclear, glutamatergic neurotransmission, calcium channels, and synaptic pruning are highly implicated (Heyes et al., 2015; Schizophrenia Working Group of the Psychiatric Genomics, 2014; Sekar et al., 2016). Our approach uncovered whole-brain activity phenotypes for genes associated with these functionalities (Figure 4A) as well as others with completely unknown functions (Figure 4B), such as *luzp2* (leucine zipper protein 2) and *gramd1b* (GRAM domain containing 1B).

**Figure 4.**
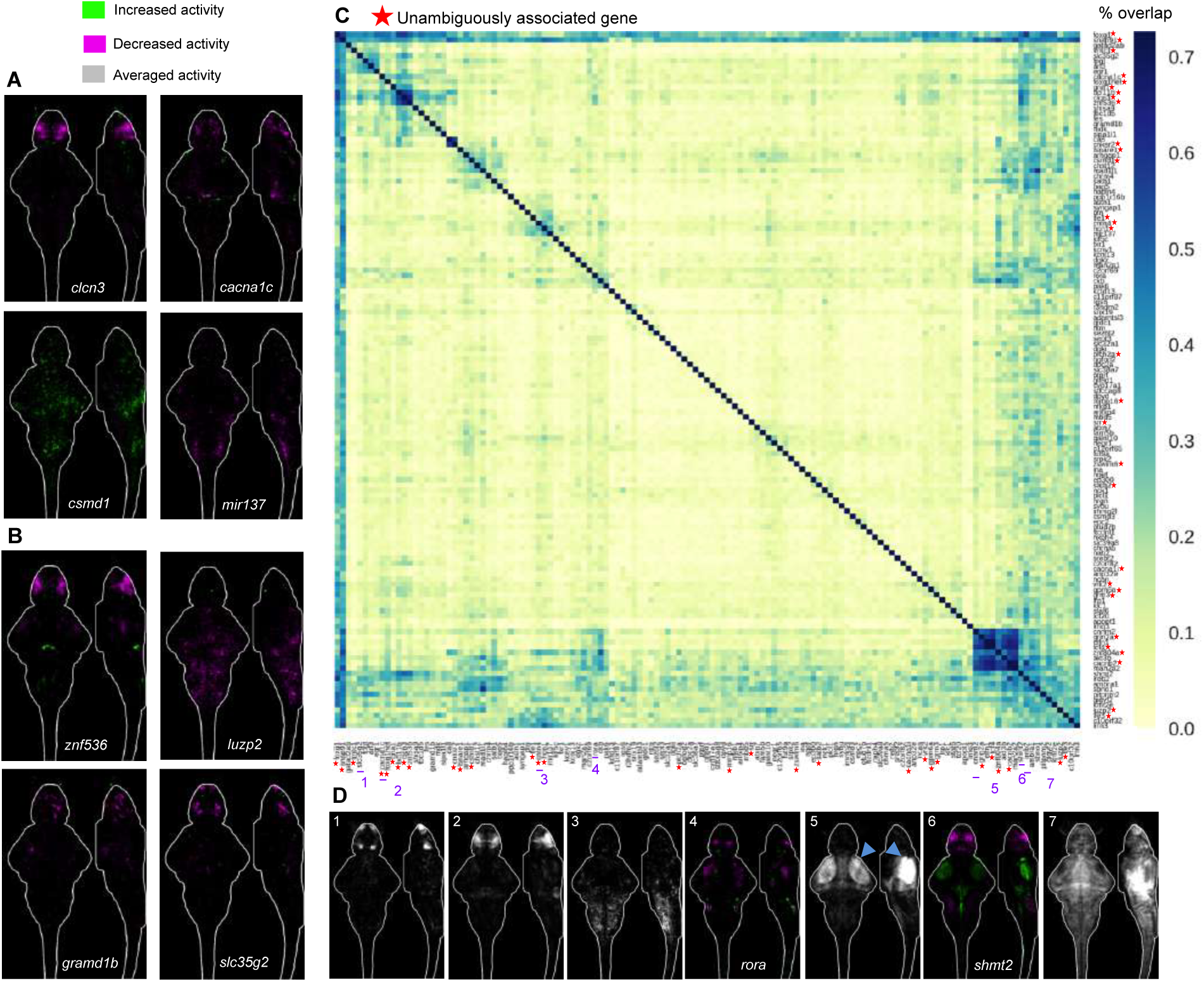
Whole-brain activity phenotypes in zebrafish mutants. (**A**) Brain activity phenotypes for genes that have been strongly implicated in schizophrenia by previous studies (Table S1). The *cacnalc* mutant phenotype shown is for heterozygous larvae because the homozygous mutant is embryonic lethal (Stainier et al., 1996). (**B**) Brain activity phenotypes for two genes that have been minimally studied and have unknown functions. (**C**) Percent overlap between mutant brain activity phenotypes was calculated between each image by comparing each brain activity signal to signal in the same location in all other mutant images. These overlaps were then sorted with hierarchical clustering using average linkage. The direction of the change in brain activity was disregarded to maximize identification of affected brain regions, and because the direction of the genetic perturbation in human patients is not clear for most genes. The numbers of image stacks that were compared to calculate significant differences in brain activity for each mutant are available in Table S2 (average N is 20 for mutant group, 28 for control group). (**D**) Sum-of-slices intensity projections, along Z- or X-axis, of significant differences between mutant and control groups of zebrafish larvae (Randlett et al., 2015). Brain activity images represent the significant differences in signal between two groups (Randlett et al., 2015), most often homozygous mutants versus heterozygous and/or wild-type siblings. See also Figure S4.

To quantitatively determine whether the same brain regions were affected across different mutants, we calculated the overlap in signal between the mutant brain activity maps (Figure 4C). Shared differences ranged from highly region-specific such as the olfactory bulb (Figure 4D, label 1) to widespread effects (Figure 4D, label 7). Hierarchical clustering of signal overlap uncovered spatial relationships between the brain areas affected in mutants (Figure 4C, Figure 4D). Unambiguously associated genes (red stars) were more likely to have overlapping signals. For example, a cluster defined by one of the two most overrepresented affected brain areas, the pallium region of the forebrain (Figure 4D, label 2), was composed entirely of unambiguously associated genes, including *bcl11b, gria1, znf536, clcn3*, and *cacna1c*. Several other mutants, such as *rora* (Figure 4D, label 4) and *shmt2* (Figure 4D, label 6), had activity changes in the pallium but were not part of the pallium cluster, since they also had changes in the other commonly affected brain region: retinal arborization field AF7 and tectum (Figure 4D, label 5). Retinal arborization field AF7 (blue arrow) is one of ten arborization fields (Robles et al., 2014) and is a visual brain area that is specifically activated during hunting (Semmelhack et al., 2014). Mutants in which this region was affected also displayed corresponding activity changes in a specific subregion of the hypothalamus, indicative of a functional connection between the two areas (Figure S5) (Filosa et al., 2016). The molecular roles of genes in the tectal cluster (Figure 4D, label 5) were diverse, ranging from transcriptional regulation (*tcf4*, *znf804a)* to glutamatergic neurotransmission (*grin2a, elfn1* (Tomioka et al., 2014)) to protein glycosylation (*man2a2*). Thus, the brain morphology and activity atlases not only uncover a large diversity of phenotypes caused by mutations in schizophrenia-associated genes but also reveal unexpected anatomical convergences despite distinct molecular functions of the mutated genes.

### Narrowing down candidates in multi-gene loci

Comparisons between phenotypes of mutants in a multi-gene locus nominated candidates for stronger consideration. Based on phenotype, we identified over 30 candidates to prioritize for future study (Table S2). For example, two schizophrenia loci (15 and 98) neighbor the genes *immp2l* and *lrrn3*, both of which were mutated in this study. The *lrrn3* gene is located within the intron of *immp2l*, making it challenging to determine the causal candidate in human genetic studies. Only *lrrn3* had strong brain activity differences (Figure 5A) and numerous behavioral abnormalities (Figure 2F), whereas loss of *immp2l* had a minimal effect on either brain activity or behavior. Our findings also added credence to candidates suggested by human studies. For example, locus 33 is located in a gene desert, and we mutated three genes (*csmd3, sybu*, and *kcnv1*) surrounding the locus. Only one of the three candidates, a second CUB and Sushi domain protein *Csmd3*, had a strong phenotype (Figure 2F, Figure 5B). This gene has a role in dendrite development and has been implicated in autism and epilepsy (Floris et al., 2008; Mizukami et al., 2016; Shimizu et al., 2003).

**Figure 5.**
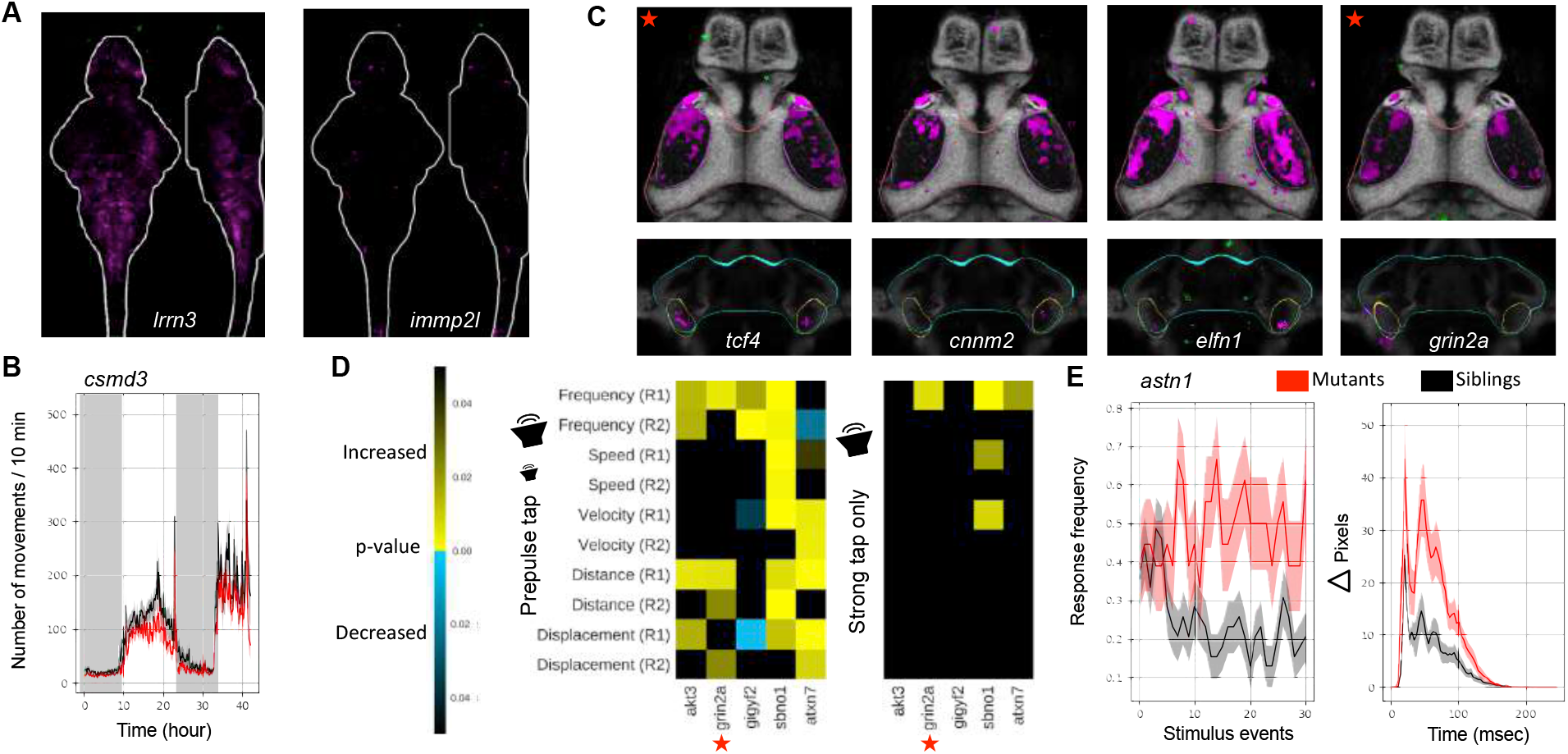
Nominating candidates in multi-gene loci by phenotype. (**A**) Brain activity data (sum-of-slices projection) for *lrrn3* and *immp2l* mutants. (**B**) Movement frequency for *csmd3* mutants. (**C**) Mutants with tectum and retinal arborization field AF7 phenotypes (representative slices). Several also have phenotypes in a small subregion of hypothalamus. See also Figure S5. (**D**) Prepulse inhibition phenotypes for five mutants. These mutant phenotypes are specific to the strong prepulse tap (Figure 2I) and do not represent a general increase in tap sensitivity to strong taps (right heatmap). Response features were calculated only on strong tap responses where the weak tap did not elicit movement. See also Figure S3. (**E**) Habituation phenotype of *astn1* mutants. Response frequency to tap events occurring every two seconds is shown in left graph, and the magnitude of responses occurring during the habituation paradigm in the right graph.

Clustering mutants by brain activity phenotype (Figure 4C) highlighted potential candidate driver genes in multi-gene loci (Table S2) when one gene in the locus shared phenotypes with unambiguously associated genes. For example, the brain activity of the magnesium transporter *cnnm2* (Arjona et al., 2014) mutant closely resembled that of the unambiguously associated mutants for the *tcf4* (Quednow et al., 2014) transcription factor and the NMDA receptor subunit *grin2a* (Figure 5C). The *cnnm2* locus is ranked 3^rd^ out of the 108 regions, and contains more than ten candidates, (Figure 1B), of which we tested five (Table S1, genepile.com/scz_gwas108). The other four mutants from this locus (*arl3, cyp17a1, c10orf32, ina*) did not have phenotypes or did not cluster with unambiguously associated genes (Figure 4C). This result nominates *cnnm2* as a likely driver of this association, highlighting the power of the large-scale approach in defining relevant genes.

Behavioral phenotypes can also help refine multi-gene loci, if behavioral differences have relevance to schizophrenia such as decreased prepulse inhibition (Swerdlow et al., 2006) and lack of acoustic habituation (Williams et al., 2013). Both of these phenotypes were observed in mutants (Figure 5D, Figure 5E, Figure 2F). Mutants with decreased prepulse inhibition included the translational repressor *gigyf2* (Kryszke et al., 2016; Morita et al., 2012), also linked to autism (Wang et al., 2016), and transcription factor *sbno1* (Takano et al., 2011), additionally identified as a possible schizophrenia candidate gene in *de novo* mutation studies (Girard et al., 2015; Girard et al., 2011). The gene *astn1*, involved in cerebellar development (Adams et al., 2002), showed a robust decrease in stimulus-driven acoustic habituation (Figure 5E). The fact that similar phenotypes are also observed in some individuals with schizophrenia (Meincke et al., 2004) nominates these genes for stronger consideration as potential drivers of the genetic association of their respective multi-gene loci.

### Mutant phenotypes of relevance to other neuropsychiatric disorders

There is substantial overlap in the risk factors for neuropsychiatric disorders (Cascella et al., 2009; Gandal et al., 2018). One condition related to schizophrenia is epilepsy. The symptoms of temporal lobe epilepsy can resemble psychosis and children with epilepsy are at an increased risk of developing schizophrenia (Cascella et al., 2009). Thus, these disorders may share common genetic underpinnings. In larval zebrafish, center avoidance has been attributed to the initiation of epileptic seizures (Baraban et al., 2005). We found sustained decreases in center time in mutants for *elfn1*, a gene involved in glutamate receptor recruitment (Tomioka et al., 2014), *hcn1*, a potassium channel, and synaptic vesicle biogenesis protein *snap91* (Fromer et al., 2016; Yao et al., 2002), among others (Figure 6A). All three of these mutants have been implicated in epilepsy in humans or mice (Koo et al., 2015; Nava et al., 2014; Tomioka et al., 2014). Correspondingly, we observed substantial increases in brain activity in these mutants, particularly in the hindbrain region for *elfn1* and *hcn1* (Figure 6B).

**Figure 6.**
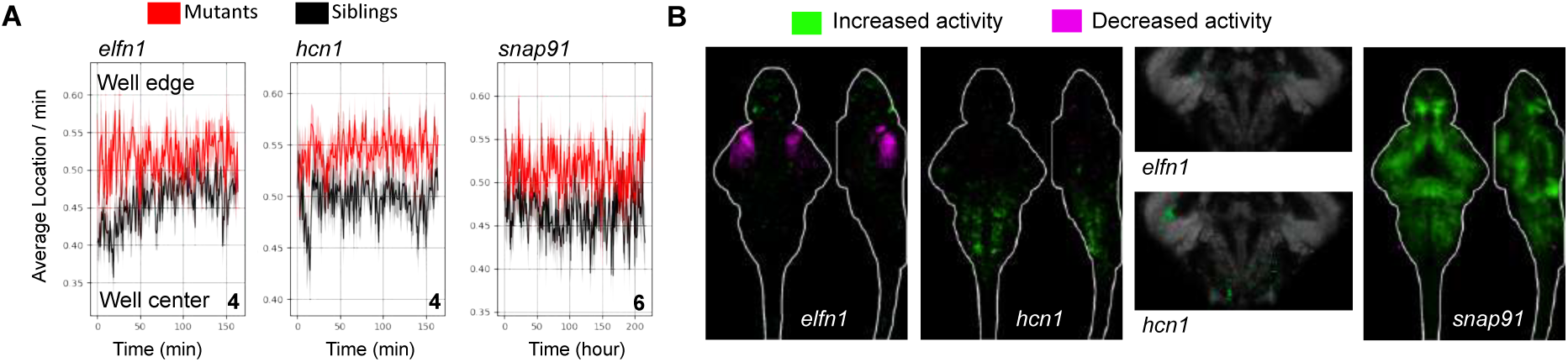
Phenotypes relevant to epilepsy. (**A**) Location preferences for three mutants, all linked to epilepsy in mammals. See also Figure S3. (**B**) Whole-brain activity (sum-of-slices projections) for the three mutants, and representative hindbrain slices for *elfn1* and *hcn1*.

A second disorder that shares genetic risk architecture with schizophrenia is autism (Gandal et al., 2018). We mutated several genes also linked to autism. One in particular, the histone methyltransferase *kmt2e* (schizophrenia locus 52 of 108), was strongly linked to autism through genome-wide association (Grove et al., 2018) and has also been observed as a *de novo* mutation (Dong et al., 2014). We found by our neuroanatomical screening that this mutant had increased brain volume (Figure 3C), which is also observed in some autistic patients (Redcay and Courchesne, 2005). This increase in brain size was unique to this gene, as the other 15 mutants with substantial morphological differences mainly had reduced volume (stackjoint.com/zbrain). The discovery of phenotypes of relevance to multiple neuropsychiatric disorders highlights the utility of this resource for catalyzing future studies in several fields.

### Neurobiological roles of top candidates

Our systematic phenotypic analyses defined top candidates and their neurobiological roles (Table S2). To explore how this dataset can direct follow-up studies, we asked whether mutant brain activity maps could yield hypotheses about potentially affected behaviors. The retinal arborization field AF7 (Figure 4D, label 5; Figure 5C) is known to specifically activate when fish hunt (Semmelhack et al., 2014), and we therefore hypothesized that hunting behavior (Jordi et al., 2015; Shimada et al., 2012) might be affected in mutants with reduced activity in this area. Indeed, we found that the mutants for the transcription factor *tcf4* (Figure 5C) showed dramatically reduced hunting behavior (Figure 7A, Figure 7B). This phenotype was highly specific, as mutants and wild type displayed similar baseline movement and visual stimulus responses (Figure 2E).

**Figure 7.**
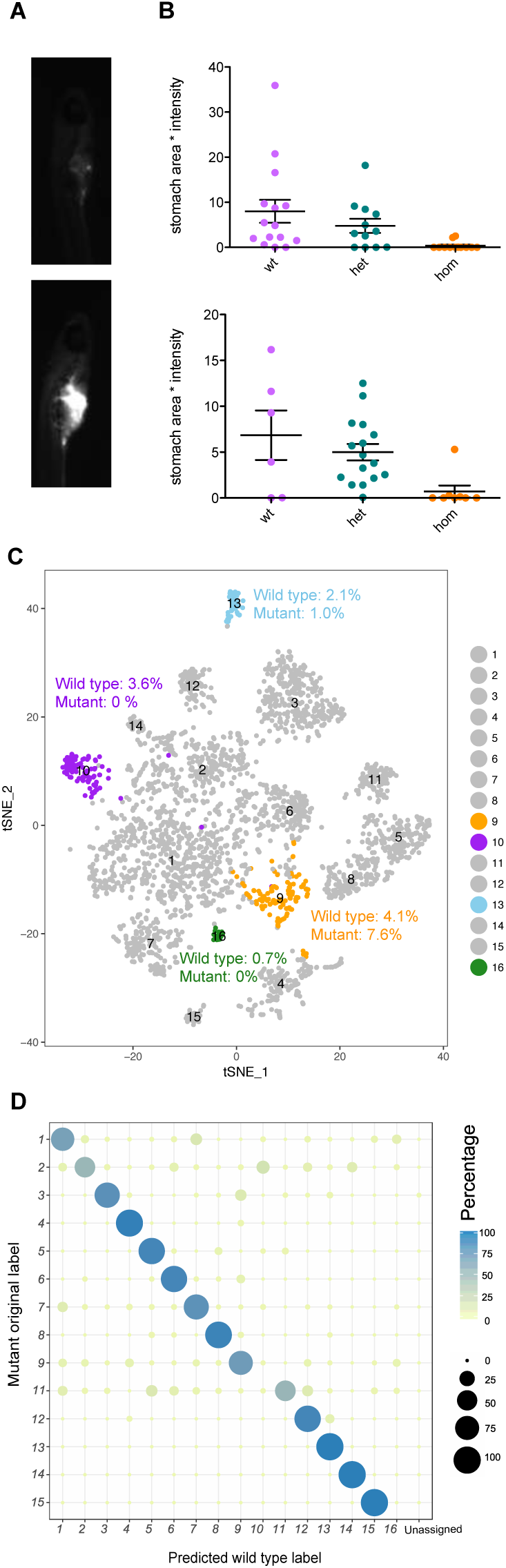
Neurobiological roles of top candidates. (**A**) Example images of larvae after hunting and consuming fluorescently labeled paramecia. (**B**) Quantification of hunting behavior by measurement of paramecia consumed. Two independent experiments are shown. (**C**) T-SNE visualization of wild type single-cell clusters obtained by clustering of 6 dpf forebrain cells. Clusters with substantial differences in mutants are highlighted in orange, purple, and blue. Cluster counts in mutant and wild type are expressed as percent of the total cell number for each sample. See also Figure S6. (**D**) Dotplot (confusion matrix) showing the proportion of cells in the mutant forebrain that were classified to wild type cluster labels. Each mutant forebrain type was assigned to a wild type cluster label if > 13% of the trees in the RF model contributed to majority vote. Proportion of cells in each row adds to 100%.

In a second test for the potential of the phenotypic database in informing follow-up studies, we studied the transcription factor *znf536* (Figure 1C). Mutants for this gene had volume loss and activity differences in the pallium region of the forebrain (Figure 3C, Figure 4B). To uncover what neuron types might be missing, we profiled the single-cell transcriptomes of dissected mutant and wild-type sibling forebrains (Figure 7C, Figure S6). This comprehensive forebrain map revealed a loss (cluster 10 and cluster 16) and reduction (cluster 12) of cell types (Figure 7C, Figure 7D). Both types uniquely express neuropeptides involved in stress and social behavior: *uts1* (Neufeld-Cohen et al., 2010) in the missing cluster 10, *penkb* (Schwarzer, 2009) in the missing cluster 16, and *tac3b* (Andero et al., 2014; Zelikowsky et al., 2018) in the reduced population (cluster 13). Corresponding with the loss of the *uts1* population was an increase in a pool of immature neurons (cluster 9), suggesting a potential developmental mechanism underlying the *znf536* mutant phenotype (Figure 7D, Figure S6). The *tac3b* population is additionally marked by *grm2a* (Figure S6), which is a potential schizophrenia therapeutic target (Griebel et al., 2016; Li et al., 2015; Mossner et al., 2008). The identification of these specifically affected types of neurons illustrates the potential of combining our phenotype database with follow-up analyses such as single-cell RNA sequencing to dissect the biological processes regulated by schizophrenia-associated genes.

## Discussion

This study establishes a zebrafish phenotypic atlas for schizophrenia-associated genes, and demonstrates the power of large-scale mutant analyses and whole-brain imaging in zebrafish in defining gene function and refining the architecture of gene-rich loci (Table S2). Our findings reinforce the contribution of genes and pathways previously implicated in schizophrenia, including glutamatergic neurotransmission (*gria1, grin2a, elfn1, clcn3*), calcium channel function (*cacna1c, cacn2b*), neurodevelopment (*bcl11b, foxg1, mir137, tcf4, gpm6a*), and complement regulation (*csmd1*).

Importantly, we uncover unexpected phenotypes for understudied genes, some of which are unambiguously associated (Table S1) with the disorder. These candidates include several with completely unknown neurobiological roles (*luzp2, gramd1b, slc35g2, znf536*). Others have undergone previous characterizations, but our analyses reveal novel phenotypes (*astn1, atxn7, cnnm2, csmd3, lrrn3, gigyf2, kmt2e, man2a2, tle3, vrk2, znf804a*). These genes implicate processes such as magnesium transport (Arjona et al., 2014) and translational control (Kryszke et al., 2016) in schizophrenia.

Our systematic approach nominates potential driver genes in multi-gene loci (*cnnm2, elfn1, shmt2, csmd3, lrrn3, akt3, gigyf2, rora, sbno1, atxn7, astn1*). This refinement is based on phenotypes shared with unambiguously associated genes and phenotypes relevant to schizophrenia pathology, such as forebrain dysfunction and prepulse inhibition defects. Notably, phenotypic commonalities (Figure 4C, Figure 5C) are detectable not at the molecular level (as in channelopathies) or cellular level (as in ciliopathies) but only at the whole-brain level. For example, sixteen seemingly unrelated genes make up the two largest groups of shared phenotypes, affecting activity in the pallium and tectum, respectively (Figure 4). It is tempting to speculate that these molecularly unrelated genes impinge on spatially or temporally overlapping developmental processes. These genes are excellent choices in gene-rich loci to be the focus of future human genetics studies.

The relevance of our dataset and pipeline extends beyond schizophrenia and zebrafish neurodevelopment. The phenotyping approach introduced in this study can be used to identify candidates and uncover mechanisms for the numerous diseases for which genome-wide association data exists (Visscher et al., 2017). Many of the genes we tested are involved not only in schizophrenia, but also in autism (Gandal et al., 2018; Grove et al., 2018), epilepsy (Cascella et al., 2009), bipolar disorder (Cross-Disorder Group of the Psychiatric Genomics, 2013; Gandal et al., 2018), depression (Wray et al., 2018), and sleep disorders (Hammerschlag et al., 2017; Lane et al., 2017). In addition to shared risk loci, some candidates have been identified by exome studies for these disorders (Dong et al., 2014; Genovese et al., 2016; Gershon et al., 2014; Girard et al., 2011; Quednow et al., 2014; Wang et al., 2016). We discovered phenotypes of relevance to both autism and epilepsy (Figure 3, Figure 6). Therefore, our established pipeline and tools can be used to screen genes that are candidates for these disorders, and future mechanistic studies of these genes will provide insights into multiple disorders. Conservation is revealed by the similarities we observe between zebrafish and mammalian phenotypes at the anatomical (*foxg1*, *bcl11b, rora, akt3* (Bergeron et al., 2017), *slc32a1*) and behavioral (*gpm6, elfn1, hcn1, snap91, akt3* (Bergeron et al., 2017)) level (Figure 4), suggesting that large-scale larval zebrafish screens complement studies in more complex systems.

The mutants described here and available to the community (stackjoint.com, genepile.com) identify among the hundreds of candidate genes at implicated loci (Schizophrenia Working Group of the Psychiatric Genomics, 2014) some three dozen genes with specific brain phenotypes (Table S2). These genes might be considered stronger candidates to be the focus of detailed mechanistic studies. For example, we discovered that the understudied transcription factor *znf536* has an essential role in development of a small subset of forebrain neurons implicated in stress and social behavior (Figure 7). As these functionalities are disrupted in patients with neuropsychiatric disease, and one of the neuron types is distinctly marked by the therapeutic target *grm2* (Griebel et al., 2016; Li et al., 2015; Mossner et al., 2008), this result is a promising lead for future study. The phenotype atlas of schizophrenia-associated genes lays the foundation for deciphering the molecular, cellular and physiological pathways underlying neuropsychiatric disease and for high-throughput screens for suppressors of mutant phenotypes (Hoffman et al., 2016; Rennekamp and Peterson, 2015; Rihel et al., 2010).

## Acknowledgements

We thank the Harvard Center for Biological Imaging for infrastructure and support, fish facility personnel Jessica Miller, Brittany Hughes, Steve Zimmerman and Karen Hurley for supporting this study, Erin Song for assistance with the hunting assay, Stephan Ripke for use of his Manhattan plot image, and William Joo, Philip Abitua, Martin Haesemeyer, Florian Engert, and members of the Schier and Engert labs for helpful feedback, and Akira Sawa for helpful comments on the manuscript. This research was supported by Damon Runyon Cancer Research Foundation (SBT), K99 MH110603 (SBT), Harvard Brain Institute Bipolar Seed Grant (AFS) and R01 HL109525 (AFS).

## Author Contributions

SBT and AFS conceived the project. SBT led all aspects of the research and did the majority of experimental work and analysis. SBT and AFS wrote the paper, with assistance from EHL, YW, SAM, and SP. LMP, EHL, CS, and JWC contributed to mutant generation and the phenotype screen. ERS, JG, SBT, EHL, SZ, and LP contributed to design and construction of the behavior rigs. SP collected and analyzed single-cell sequencing data, with assistance from SBT. YW designed and implemented the image clustering, and provided input on the behavior analyses. OR shared phosphorylated-Erk protocols prior to publication and provided helpful feedback, including assistance in establishing the DBM analysis. SAM provided valuable input on schizophrenia genetics. NSM worked with SBT to design and build stackjoint.com and genepile.com.

## STAR Methods

### CONTACT FOR REAGENT AND RESOURCE SHARING

Further information and requests for resources and reagents should be directed to and will be fulfilled by the Lead Contact, Summer B. Thyme.

### EXPERIMENTAL MODEL AND SUBJECT DETAILS

#### Zebrafish husbandry

All protocols and procedures involving zebrafish were approved by the Harvard University/Faculty of Arts & Sciences Standing Committee on the Use of Animals in Research and Teaching (IACUC; Protocol #25-08). Mutants were generated by simultaneous injection of one to four guide RNAs (gRNAs), made as previously described (Thyme et al., 2016), and approximately 0.5 nL of 50 μΜ Cas9 protein into EK*/nacre (mitfa^−/−^*) (Lister et al., 1999) embryos. Sequences of all gRNAs and resulting mutations are available on genepile.com. The resulting mosaic adults with a germline mutagenic deletion were crossed to either EK or *nacre* and the offspring were mated to each other to generate homozygous mutants. If the human gene of interest was duplicated in zebrafish, homozygous mutations were generated for both orthologues. Tests of duplicated genes were most often completed with double homozygous (−/−:−/−) compared to +/−;+/−, +/−;−/−; and −/−:+/− siblings (exact comparisons described in Table S2). All larvae were genotyped (Truett et al., 2000) to generate lines and for all phenotypic characterizations. Zebrafish were raised at 28.5°C on a 14/10 hour light/dark cycle. Behavioral experiments were conducted on the same cycle and at the same temperature. Fish used for all assays were grown in 150 mm petri dishes in fish water containing methylene blue, at a density of less than 160 larvae per dish, and debris was removed twice prior to 4 dpf.

### METHOD DETAILS

#### Brain activity and morphology

Phosphorylated-Erk antibody staining was conducted as previously described (Randlett et al., 2015). Larvae were fed paramecia in the morning of 5 dpf. On 6 dpf they were maintained in a quiet environment and fixed in the mid-afternoon in 4% paraformaldehyde (PFA) (Supplier Polysciences) diluted in 1X Phosphate Buffered Saline with 0.25% Triton (PBST). To ensure rapid fixation, as Erk is phosphorylated in neurons 10 seconds after calcium signaling (Randlett et al., 2015), larvae were poured into a mesh sieve and immediately submerged in fix. After overnight fixation, larvae were washed three times for 15 minutes each in PBST. Pigment on larvae was bleached with a 1% H_2_O_2_/3% KOH solution, followed by three 15-minute washes in PBST. Larvae were then incubated at 70°C for 20 minutes in 150 mM TrisHCl pH 9.0, to retrieve antigens (Inoue and Wittbrodt, 2011), followed by two five-minute washes in PBST. Permeabilization of tissue was conducted by incubation in 0.05% Trypsin-EDTA on ice for 45 minutes. After three 15-minute washes and blocking for one hour (PBST with 2% normal goat serum (NGS), 1% bovine serum albumin (BSA), and 1% dimethyl sulfoxide (DMSO)), primary antibodies were diluted in blocking solution and incubated overnight. The phosphorylated Erk antibody (Cell Signaling, #4370) and total Erk antibody (Cell Signaling, #4696) were diluted both 1:500. The following day larvae were rinsed three times for 15 minutes each with PBST, and alexa-fluorophore conjugated secondary antibodies (Life Technologies) were diluted 1:500 in blocking solution for a second overnight incubation. Following three 15-minute PBST washes, larvae were mounted in 2% low-melting agarose in 1X PBS. Images were collected using either Zeiss LSM 780 or 880 upright confocal microscopes with a 20X/1.0 NA water-dipping objective. Of the 132 mutants, 78 were independently repeated to confirm results (Table S2, stackjoint.com/zbrain). Larvae were removed from the agarose and genotyped after imaging.

#### Behavior

The majority of assays were conducted with between 24 and 96 larvae in a multi-well plate (exact Ns in Table S2). Fish were transferred to 96-well square 0.7 mL plates (E&K Scientific) on the afternoon of 4 dpf. Plates were filled with standard fish water containing methylene blue, transferred to ice until locomotion ceased, and sealed with optical adhesive films (ThermoFisher Scientific). Experiments were conducted at approximately 28°C. Larval locations were monitored at approximately 30 frames-per-second (fps) with 1088 × 660-pixel resolution during the entire time course using a Grasshopper3 camera (FLIR). To monitor stimulus responses, the camera generated one-second long movies at 285 fps. Tap stimuli were delivered by computer-controlled increases in voltage to a solenoid attached to the apparatus. Tap strengths for weak and strong taps were determined based on percent of larvae responding. The strong was set as high as possible, with almost 100% of the fish responding if they had not habituated to the stimulus. The weak tap was set so that less than 10% of the larvae responded. We repeated behavioral assessments for 73 of the mutants, including 46 mutants of the 49 designated as having a behavioral phenotype (Figure 2F, Table S2). The protocol lasted two days and was analyzed as 14 time windows, some of which contained stimuli that are detailed as follows:

*Time windows:*

1. Night 1, 11:59 pm - 8:59 am, 9 hours, no events and lights were off.
2. Morning transition 1, 9:00 am - 9:10 am, 10 minutes, lights turned on to a low level at 9:00 am.
3. Morning light flash, 9:10 am - 11:15 am, 125 minutes, twenty 1-second long light flashes with 1-minute spacing starting at 9:11 am.
4. Day baseline, 11:15 am - 2:00 pm, 165 minutes, no events and lights were at a standard intermediate level.
5. Day taps, 2:00 pm - 7:22 pm, prepulse inhibition and short-term tap habituation, 5 hours and 22 minutes. Randomly intermixed strong, weak, and prepulse taps occurred from 2:00 pm - 5:00 pm, with spacing of 30 seconds. Three blocks of tap habituation occurred from 5:30 pm - 7:07 pm. Each block consists of 10 strong taps with spacing of 30 seconds (pre-habituation), followed by a 10-minute break and then 33 strong habituating taps with 2 second spacing. Of the 36 taps, numbers 4, 19, 36 are high-speed (computer cannot process high-speed movies every 2 seconds) and used to calculate habituation phenotypes This habituation section was followed by a 3-minute break, and then 10 taps separated by 30 seconds. Then the second habituation block began (taps every 2 seconds), and the cycle was repeated once more for the 3^rd^ block.
6. Evening, 7:22 pm - 10:59 pm, 217 minutes, no events and lights were at standard intermediate level.
7. Night transition, 11:00 pm - 11:10 pm, 10 minutes, lights turned off at 11:00 pm.
8. Night 2, 11:10 pm - 1:00 am, 110 minutes, no events and lights were off.
9. Night stimulation, 1:00 am - 7:00 am, including tap habituation, prepulse tests, tap response threshold tests, and light flash stimulation, 6 hours. This window included 1 hour of random strong, weak, and prepulse taps, followed by one habituation block, followed by twenty light flashes.
10. Night 2b, 7:00 am - 8:59 am, second section, 119 minutes, no events and lights were off.
11. Morning transition 2, 9:00 am - 9:10 am, 10 minutes, lights turned to standard intermediate level at 9:00 am.
12. Morning 2, 9:10 am - 10:00 am, 50 minutes, no events and lights were at standard intermediate level.
13. Dark flashes, 10:00 am - 5:00 pm, 7 hours, four 1-hour blocks of 1-second long dark flashes with 1 minute spacing. Each block is spaced by 1-hour of rest. The first 10 and last 10 flashes of each of the four blocks are recorded at high-speed and analyzed to determine dark flash responsiveness over the course the experiment (flash responses 11-50 are not analyzed).
14. Heat stressor, after 5:00 pm, >70 minutes, manually turned on water heater and system took 10-15 minutes to reach a 10°C increase that was maintained for approximately 1 hour.

#### Fluorescent *in situ* hybridization

Fluorescent RNA ISHs were performed as previously described (Pandey et al., 2018; Ronneberger et al., 2012) using 6 dpf *mitfa−/−* larvae that were either wild type (Figure S4) or a blinded mix of *znf536* mutant larvae and control siblings (genotyped after imaging). To generate probes, gene fragments were amplified from with Phusion polymerase (New England Biolabs, M0530L). Sequences for data shown in Figure S4 are available on genepile.com. The fragments were cloned, and resulting plasmids were linearized with restriction digestion and purified using a PCR clean up kit (Omega Cycle Pure Kit). Digoxigenin-labeled RNA probes were generated using the RNA labeling kit (Roche) and purified using Total RNA clean up kit (Omega, R6834). The quality of the probes was confirmed on an agarose gel and concentration was quantified using Nanodrop. The final product was diluted to 50 ng/μL in Hybridization Mix (HM+) buffer (50% formamide, 5X Saline Sodium Citrate (SSC) buffer, 0.1% Tween 20, 5 50 μg/mL heparin, mg/mL torula RNA).

Larvae were fixed in 4% PFA diluted in PBS at 4°C overnight, and subsequently rinsed three times in *in situ* PBST (PBS with 0.1% Tween 20). Larvae were then dehydrated with increasing methanol (MEOH) concentration diluted in PBST, incubating for 10 minutes each in 25%, 50%, 75%, 100%, and 100% MEOH. Following at least one overnight at −20°C, larvae were rehydrated with decreasing concentrations of MEOH diluted in PBST (10 minutes incubation, 75%, 50%, 25%, 0% four times). Larvae were then incubated in Proteinase K for 1 hour at a concentration of 10 μg/mL and the reaction was halted by incubating in 4% PFA for 20 minutes. Larvae were pre-hybridized in HM+ buffer for 2 hours at 65°C. Probes, normalized to 3.33 ng/μL, were denatured at 70°C for 10 min before hybridization, which was conducted overnight in HM+ buffer at 65°C. Larvae were washed the next day at 65°C first for 20 minutes each in HM+ buffer, 75% formamide diluted in 25% SSCT buffer (2X SSC with 0.1% Tween 20), 50% formamide with 50% SSCT, 25% formamide with 75% SSCT, 100% SSCT, 100% SSCT, and then three times for 30 minutes in 0.2X SSCT and twice in room temperature for 10 minutes in TNT (100mM Tris-HCl, pH 7.5, 150mM NaCl, 0.5% Tween 20). Blocking for a minimum of 1 hour (TNTB: 1% blocking agent in TNT) at room temperature followed, and then larvae were incubated with peroxidase-conjugated anti-digoxigenin-POD antibody (Roche 11 207 733 910) at 4°C overnight using a 1:400 dilution in TNTB. The next day, larvae were washed 8 times for 15 minutes each in TNT, stained for 1 hour in the darkness (Perkin Elmer TSA Plus Cyanine 3 System, NEL744001KT), and washed three times for 5 minutes each in TNT.

#### Hunting assay

Paramecia were stained as previously described (Jordi et al., 2015; Shimada et al., 2012), but using the 1,1′-Dioctadecyl-3,3,3′,3′-Tetramethylindodicarbocyanine, 4-Chlorobenzenesulfonate Salt (DiD’) dye (ThermoFisher Scientific D7757). Cultured paramecia (approximately 100 mL) were purified by filtration through a fine mesh that retains them (pore size 20 μΜ), concentrated by centrifugation for 5 minutes at 3,000 rpm, and resuspended in 800 μL of water. The solid dye was dissolved in ethanol to make a 2.5 mg/mL working solution, of which 4 μL was added to the tube of paramecia. Paramecia were stained for 2 hours with gentle rocking, spun for 5 minutes at 3,000 rpm, washed two times with water, and resuspended in 1 mL of water. The food was used at approximately 100 μL for ten larvae, and the larvae in a 150 mm petri dish were allowed to feed for 15 minutes before they were fixed in 4% PFA in PBS overnight. Prior to the assay, larvae had not eaten for over 12 hours. The next morning, larvae were rinsed three times in PBS and their stomach intensity was imaged on a Zeiss Axio Zoom.V16 Stereo Microscope with a Cy50 filter. Quantified was completed using the 3D objects counter in ImageJ with a constant threshold. All larvae are tested together and genotyped after imaging of stomach labeling.

#### Cell isolation and library preparation for 10X scRNA-seq

Cell isolation and capture in droplets was performed as previously described (Pandey et al., 2018) with the following minor modifications. Eight 6 dpf larval forebrains were dissected for each sample (wild type and *znf536* homozygous mutant) in Neurobasal (ThermoFisher Scientific 21103049) supplemented with 1x B-27 (ThermoFisher Scientific 17504044) within approximately forty minutes. Forebrains were incubated in 20 units/mL papain (Worthington Biochemical Corporation LK003150) for 12 minutes at 37°C, followed by gentle trituration approximately 20 times and spinning at 300xg for 5 minutes. Dissociation was halted by resuspension in 1 mL of 1.1 mg/mL papain inhibitor in Earle’s Balanced Salt Solution (EBSS), spun at 300xg for 5 minutes. The resulting pellet was then washed in Neurobasal supplemented with B27 before final resuspension in 50 μL PBS + 200 μg/mL BSA. To avoid cell loss, during each wash, the supernatant was saved and spun a second time to recover cells. These cells were then mixed with the final cell suspension to increase cell count. After confirming a cell viability of greater than 80% using trypan blue, cells were counted on a hemocytometer and loaded onto the 10X Chromium system at a concentration of approximately 150 cells/μL. Libraries were prepared according to the manufacturer’s instructions.

### QUANTIFICATION AND STATISTICAL ANALYSIS

#### Brain activity and morphology analysis

Images were registered to a standard zebrafish reference brain using Computational Morphometry Toolkit (CMTK) (Jefferis et al., 2007; Rohlfing and Maurer, 2003). All phosphorylated-Erk images were normalized with a total-Erk stain. The Mann-Whitney U statistic Z score is calculated for each voxel, comparing between the mutant and control groups using MapMAPPING (Randlett et al., 2015). The significance threshold was set based on a false discovery rate (FDR) where 0.05% of control pixels would be called as significant. Structural analysis with CMTK was based on deformation-based morphometry (Cachero et al., 2010; Rohlfing and Maurer, 2003) to localize changes in brain volume, and the differences were calculated the same way as for the brain activity using MapMAPPING but with a 0.005% FDR threshold. When multiple comparisons were made, such as in the case of duplicated genes, the comparison with the most extreme phenotype was selected to represent the gene. The numbers of mutant and control fish in each experiment are available in Table S2 and on stackjoint.com/zbrain. All image processing and computational analyses were completed on the Harvard Odyssey cluster.

#### Image similarity analysis

To cluster the mutants based on their neural activities, the following steps were taken:

1. Calculated the absolute difference in pixel intensities between each mutant and its wild-type / heterozygous sibling control, as a measurement of neural activity changes in the mutant.
2. Set the pixels outside the brain region to zero intensity.
3. Defined each pixel with an intensity greater than 50 as a *signal* of activity change. The threshold of 50 was chosen such that background intensities were best separated from real signals based on test images.
4. If a brain activity imaging experiment was independently repeated (78 of the 132 mutants, 59 of those designated as having brain activity phenotypes), irreproducible signals were eliminated. First, the brains were segmented into four major brain regions: telencephalon, mesencephalon, rhombencephalon, and diencephalon (Randlett et al., 2015). Signals in a brain region were discarded if they occupied very few pixels (<1/5000 of the total number of pixels in that brain region), or if they occupied pixels only in one of the two images. The pixels were also discarded if the directionality of signals (increased or decreased) was not the same in both images. After the elimination of irreproducible signals, the two images were averaged to obtain one image for each mutant. These images were used for calculating pixel sums (Figure 1G, Figure 1H, Figure 4C).
5. Dilated each signal in 3D by 8-pixels in diameter in x and y dimensions, and 4-pixels in z dimension to define a *region* of activity change for each mutant (Equation 1).
6. For each pair of mutants, determined the *overlap* region of activity changes (Equation 3). Then, calculated the percentage of signals within the overlap region for each mutant and averaged the two percentages to obtain the similarity score between the two mutants (Equation 3).
7. Repeated step 6 for all mutant pairs to construct a similarity matrix, which was then used as the distance matrix for a hierarchical clustering algorithm.

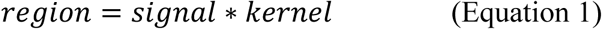

where *signal* is the binary 3D matrix that marks pixels with activity change with 1, * indicates convolution operation, and kernel is defined as:

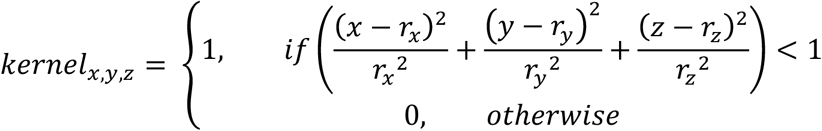

where rx, ry, and rz are the radius for dilation along the 3 dimensions.

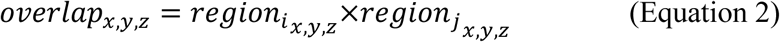

where *region_i_*, and *region_j_* are regions of activity change for the i-th and j-th brain respectively, and x, y and z represent indices in the 3D matrices. The resulting *overlap* is a 3D matrix with 1 marking the overlapping pixels.

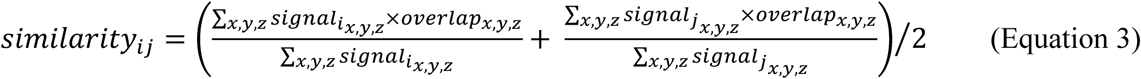

#### Behavior Analysis

To determine whether mutants had a significant difference in either frequency of movement, features of movement, location preferences, or response to a stimulus, p-values for individual metrics were merged (eg, velocity and distance of movements are both considered features) as described in Figure S2B. For data with a time component, a linear mixed effects model was used to calculate significance:

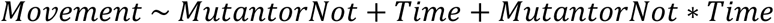

The model assumes that the movement of the fish can be approximated as a linear combination of (1) a constant (baseline) movement level, (2) a trend of movement in time, (3) a change in the baseline movement level due to the mutation, and (4) a change in the trend of movement due to the mutation. *MutantorNot* was the parameter evaluated for significance. The advantage of the linear mixed effects model is that it takes into account the longitudinal order of the data points, and therefore has the potential of recognizing consistent changes within a limited time window that might be missed if the order is disregarded. Kruskal-Wallis one-way ANOVA was used to compare stimulus responses for mutant and control groups based on the mean response metric for each fish. Once significance was determined for individual metrics (described below), the p-values were merged into 71 categories or behavior assays (Figure S2). The merging was necessary to diminish false-positive findings. The merging approach was empirically defined as giving a <10% false discovery with randomized batches of wild-type larvae (Figure S2B).

##### Behavior metrics

For all average measures, the metric is averaged for each fish prior to significance comparisons. An increased or decreased mutant response (blue or yellow in heatmaps, Figure 2) in features of movement or stimulus responses depends on what metric is most significantly different. It is feasible that a mutant could have decreased speed and increased distance, or increased frequency with decreased distance in the case of a stimulus response.

*Movement frequency metrics* (12): Number of bouts (based on distance) / min, Number of bouts (based on distance) / 10 min, Number of bouts (based on delta pixels) / min, Number of bouts (based on delta pixels) / 10 min, Active seconds / min (based on distance), Active minutes / 10 min (based on distance), Active seconds / min (based on delta pixels), Active minutes / 10 min (based on delta pixels), Average interbout interval (seconds) / min (based on distance), Average interbout interval (seconds) / 10 min (based on distance), Average interbout interval (seconds) / min (based on delta pixels), Average interbout interval (seconds) / 10 min (based on delta pixels). A “bout” is a movement that passes certain threshold parameters, and it can be calculated based on either change in pixels between frames (threshold of 3 frames and 3 pixels) or based on actual distance moved in pixels (threshold of 2 frames and 0.9 pixels). The “interbout” is the time between movements.

*Movement features metrics* (14): Average bout cumulative delta pixels (pixels) / min, Average bout cumulative delta pixels (pixels) / 10 min, Average bout distance (pixels) / min, Average bout distance (pixels) / 10 min, Average bout displacement (pixels) / min, Average bout displacement (pixels) / 10 min, Average bout time (milliseconds) / min (based on distance), Average bout time (milliseconds) / 10 min (based on distance), Average bout time (milliseconds) / min (based on delta pixels), Average bout time (milliseconds) / 10 min (based on delta pixels), Average bout speed (pixels / millisecond) / min, Average bout speed (pixels / millisecond) / 10 min. Average bout velocity (pixels / millisecond) / min, Average bout velocity (pixels / millisecond) / 10 min.

*Location in well metrics* (8): Fraction of bout time in well center / min, Fraction of bout time in well center / 10 min, Fraction of interbout time in well center / min, Fraction of interbout time in well center / 10 min, Average bout rho / maximum rho / min, Average bout rho / maximum rho / 10 min, Average interbout rho / maximum rho / min, Average interbout rho / maximum rho / 10 min. The value rho is the radius in polar coordinates, used to determine location in well.

*Stimulus metrics* (15-148, depending on number of sections combined, e.g. weak taps occur before the strong tap in prepulse and also independently, measures for both are merged): Frequency, Latency (milliseconds), Displacement (pixels), Cumulative distance (pixels), Area moved (pixels) (Figure 2D), Time (milliseconds), Speed (pixels / millisecond), Velocity (pixels / millisecond), Cumulative delta pixels, Peak change in delta pixel per response (pixels), Peak speed in per response (pixels / millisecond), Maximum change in delta pixels from average response trace (delta pixels in every frame), Location of maximum change in delta pixels (millisecond), Maximum change in distance from average trace (distance in every frame), Location of maximum change in distance (millisecond). In addition to all the parameters described, all tap responses were further analyzed based on the same parameters for two subsets of responses: canonical escapes and weaker responses that follow if the larva did not escape. “Big” responses (e.g. true escapes) were designated as having a velocity of >0.2 pixels / millisecond and having a latency of less than 25 millisecond from the start of recording (not from the actual tap occurrence). These parameters were determined by analysis of responses to the strong tap, which are true escapes the vast majority of the time. The “small” responses were every other response that occurred up to 75 milliseconds after the start of recording. All frequency phenotypes for prepulse inhibition experiments shown in Figure 5D represent changes in frequencies for “big” responses (true escapes).

#### Analysis of single-cell sequencing data

First, cells were filtered to remove those that contained less than 200 genes and those in which > 6 of transcript counts were derived mitochondrial-encoded genes. Similarly, genes detected in less than 5 cells were removed. All cells derived from regions other than the forebrain, such as habenula and olfactory bulb, were removed. Similarly, non-neuronal types such as microglia and vascular cells were also removed from further analysis. The residual matrix was then scaled, centered and used for further analysis.

To select highly variable genes, we used a combination of a UMI based method described recently (Pandey et al., 2018) and Seurat’s (Satija et al., 2015) variable gene selection approach. The resulting expression matrix across the highly variable genes was then used to perform dimensionality reduction and clustering using Seurat.

To compare signatures between mutant and wild type datasets, both datasets were independently clustered as described. We used a multi-class Random Forest Classifier exactly as described in (Pandey et al., 2018). Briefly, the classifier was built on the most variable genes across both datasets using 1000 trees with R package RandomForest. The classifier was trained on 70% of the wild type dataset and tested on the remaining 30% of the cells (Figure S7C). Each cell in test set was only assigned into a label if a minimum of 13% of the trees in the forest converged onto a decision. Otherwise the cells were unassigned. The resulting classifier was then used to predict wild type labels for the cells in the mutant dataset.

### DATA AND SOFTWARE AVAILABILITY

All mutants have been cryogenically preserved as sperm, and are available upon request. All raw behavior and imaging data are available upon request. The 10X raw sequencing data has been deposited in GEO under the codes GSE115427. Labview code for tracking larvae and generating high-speed movies is available from the Harvard CBS Neuroengineering core at https://github.com/cbs-ntcore/Schier-Lab. Code for analyzing high-speed movies, processing both slow- and high-speed tracking data to generate behavior graphs, and for clustering brain activity maps are available at https://github.com/sthyme/ZFSchizophrenia. Code for calculating larvae locations and changes in pixels per frame (delta pixels) in high-speed stimulus response movies is partly based on previously published python tracking code (Conklin et al., 2015).

### ADDITIONAL RESOURCES

All mutant allele information is available on the following website: genepile.com/scz_gwas_108

Brain activity maps for mutants are available on the following website: stackjoint.com

stackjoint.com is a community resource, where users can upload their own data and share it with a limited set of colleagues or make it publicly available. Images can be hosted from any location chosen by the user. Any stack of images (any size, resolution, or number) can be shared through this site on stackjoint.com/basic, and zebrafish data that is registered to the Z-Brain can add to a growing repository of 6 dpf zebrafish neuroimaging data on stackjoint.com/zbrain. Users can upload and analyze their own zebrafish brain activity maps to identify regions of altered signal, based on Z-Brain anatomical masks available in this online resource. Colors of differences maps (activity, structure, or any antibody stain of interest) are designated as green for increased signal in test group (gene mutant, drug, or stimuli) and magenta for decreased. The current map naming convention for mutant data is

*Gene_AnalysisType_NofComparedAnimals_ExperimentalInfo_RunNumber_SignificanceCutoff e.g., ambra1_structure_13homhomover31homhet_hethomfxhomhetm_2nd_p00005*

where analysis type could be any antibody name, drug or stimuli type could replace gene name, and experimental information could be anything of relevance such as concentration of a drug. If uploading large amounts of data to this site, please contact for assistance in streamlining the process.

